# Benefits of dual-tasking on implicit sensorimotor adaptation

**DOI:** 10.1101/2025.06.18.660484

**Authors:** Benjamin Miller-Mills, Tara Kwan, Timothy J Carroll, Eugene Poh

## Abstract

Attention plays a crucial role in maintaining precision and effectiveness in goal-directed actions. Although there is evidence that dividing attention across tasks impairs performance in various domains, the impact of attention on sensorimotor adaptation remains inconclusive, with some studies reporting deficits and others showing no effects. Because sensorimotor adaptation arises from the interaction of explicit and implicit processes, this discrepancy may reflect differential effects of attention on each process. Here, we investigate how divided attention influences implicit sensorimotor adaptation using an error-clamp paradigm, coupled with a random dot kinematogram (RDK) motion coherence discrimination task. We also assessed whether the timing of the secondary task affects error processing during sensorimotor adaptation by presenting the RDK either during the outward movement (coinciding with error feedback), or the inward movement (following error feedback). We observed that attentional manipulation influenced implicit sensorimotor adaptation only when the RDK was presented on the outward movement, not the inward movement. Remarkably, implicit sensorimotor adaptation was enhanced when attention was divided, compared to when attention was focused entirely on the adaptation task. This suggests that implicit sensorimotor adaptation is sensitive to attentional demand, particularly during the time window where error feedback is received.

## INTRODUCTION

Sensorimotor adaptation is a fundamental aspect of human behavior, encompassing the capacity to adjust and refine movements in response to changing environmental conditions or internal states. This adaptive process is vital for maintaining precision and effectiveness in goal-directed actions, whether in complex skill execution or everyday tasks (1, 2). Because the sensory-motor interactions that support goal-directed actions require representations that span the body and the environment (3–6), this adaptive behavior often occurs while our cognitive resources and attention are divided between monitoring the internal state of our body, and the external states of behaviorally relevant environmental stimuli. For instance, when driving a car, individuals must adjust body movements like steering and pedal control to meet the varying demands of the road while simultaneously allocating attention to external factors such as avoiding pedestrians and signaling turns. Because attention has been viewed as a necessary resource that supports cognitive function, it has been frequently demonstrated that dividing attention across multiple tasks is detrimental to performance in a variety of motor tasks (7, 8). However, the literature paints an inconsistent picture as to how dividing attention impacts sensorimotor adaptation.

A common approach to assess the relationship between attention and sensorimotor adaptation has been to present a concurrent attention dividing task while participants perform a sensorimotor adaptation task. The adaptation in such dual-task conditions is then compared to single-task conditions in which only the sensorimotor adaptation task was performed. This dual-task methodology has been used extensively across various sensorimotor adaptation tasks - including prism adaptation (9–11), visuomotor adaptation (12–14) and force field adaptation (15, 16). In most of these studies, sensorimotor adaptation was impaired when participants engaged in a concurrent, secondary task. For example, Taylor and Thoroughman (16) observed that performing a force field adaptation task concurrently with an attention dividing semantic categorization task interfered with the ability to adapt to the changed dynamic conditions of the reaching movement. Specifically, increasing the difficulty of the semantic categorization task led to greater impairment in compensation for movement errors on the following trial. A separate study by the same authors found that the impairment in force field adaptation was most pronounced when the attention-dividing secondary task (this time a frequency discrimination task) temporally coincided with the movement error created by forces being applied to the hand (15). Furthermore, in a prism adaptation task, the introduction of a secondary mental arithmetic task led to increased movement variability and a reduced rate of adaptation (10). In general, these results are consistent with the traditional notion that dividing attention with a secondary task will have detrimental effects on sensorimotor adaptation. Moreover, the temporal specificity of the secondary task suggests a mechanism of interference related to error encoding.

In contrast, other studies have observed that dividing attention with a secondary task did not impair sensorimotor adaptation (17–21). For example, when various attention dividing secondary tasks (e.g., rapid visual serial presentation, brightness discrimination, and auditory frequency discrimination) were paired with a primary visuomotor adaptation task (45° visuomotor rotation), Song & Bedard (20) showed that dividing attention during the secondary task did not affect the rate of adaptation, irrespective of secondary task requirements or the sensory modality, compared to a group that performed the visuomotor adaptation task alone. Intriguingly, in a subsequent recall phase, the authors found that the visuomotor adaptation acquired concurrently with a secondary task was only recalled under the same dual-task conditions. When examined in the absence of the secondary task, there was a substantial reduction in aftereffects for the sensorimotor adaptation, with performance reverting to levels observed in naïve participants (20). This result suggests that dividing attention during dual-task performance does not always impair sensorimotor adaptation. Instead, it is the consistency of the attentional state that serves as a context to enable the retrieval of the sensorimotor memory.

These diverging effects of divided attention on sensorimotor adaptation may reflect underlying differences in the mechanisms that drive sensorimotor adaptation. Current theories propose that sensorimotor adaptation arises from a combination of explicit and implicit processes (22–26), each of which may be differentially influenced by varying attentional demands. For example, when a visuomotor rotation (23) or force-field perturbation (24) is introduced abruptly, participants can become conscious of the perturbation and develop explicit re-aiming strategies to improve performance. Such re-aiming strategies require the recruitment of cognitive resources to develop explicit knowledge about the perturbation (23, 27). Therefore, dividing attention via secondary task (e.g., re-aiming), should impair explicit processes during sensorimotor adaptation. In contrast, implicit processes are thought to be mainly driven by sensory prediction errors - defined as the difference between predicted and observed sensory outcomes during motor performance (28). It has been suggested that this implicit learning component does not engage attentional resources in general (27, 29) and is therefore more likely to remain unaffected by an attention- dividing secondary task. It is thus conceivable that the differing effects of dual-tasking reported to date may have been driven by differing contributions from explicit and implicit processes during various sensorimotor adaptation tasks.

Previous studies have attempted to shed light on this issue using indirect measures of explicit and implicit components during sensorimotor adaptation. However, they have reported little influence of attention for either process (13, 30). For example, in Langsdorf et al. (30), after learning a visuomotor rotation with a concurrent 1-back memory recall task, implicit sensorimotor adaptation was inferred from the size of the aftereffect, and explicit contributions were inferred through reports of strategic re-aiming. Intriguingly, while they observed impaired performance during the visuomotor rotation during dual-task conditions, there was no effect of dual-task interference found during the explicit or implicit posttest measures. This suggests that dividing attention with a secondary task may disrupt motor performance, but not necessarily the explicit and implicit components of sensorimotor adaptation measured via post-perturbation tests (i.e., motor memory).

Another indirect method of examining the potential contributions of explicit and implicit processes during sensorimotor adaptation is to compare responses to both gradual and abrupt perturbations (31). Using this approach, Galea et al. (13) found that introducing a secondary attention-dividing (vocal shadowing) task impaired adaptation to both abrupt and gradual visual displacements by a similar degree. This is intriguing, as abrupt perturbations were previously thought to depend more on high-level explicit strategies, presumably making them more vulnerable to attentional manipulations (31). However, it has been suggested that awareness of visuomotor rotations does not always lead to the use of explicit strategies, particularly when the perturbation is small (32, 33). Taken together, because the additional attentional demands of secondary tasks have been previously imposed when both explicit and implicit processes might be acting concurrently, and attentional effects have been inferred indirectly from post-test measures, the influence of attention on these distinct adaptive processes remains unclear.

Beyond the dichotomy of explicit and implicit processes, the temporal specificity of a secondary task may also modulate the processing of movement errors during sensorimotor adaptation (15). For example, in the studies conducted by Song and colleagues (17–21), the attention-dividing secondary task was presented during the movement for a total duration of 1.5s, spanning both the outward movement stage (reaching towards the target), and the inward movement stage (returning to the starting position). Therefore, it is conceivable that the observed absence of dual-task interference may reflect the lack of temporal specificity between the timing of the movement error, and performance of the secondary task (15). Interestingly, Stadler (34) has suggested that an attentionally demanding task may impair sensorimotor adaptation, not because it divides attention, but rather because it disrupts the normal organization of successive events in a trial. Under this lens, it is critical to investigate the temporal coincidence of movement error and secondary task presentation; ideally with the secondary task being temporally tied to either the outward movement or the inward movement exclusively during the performance of the sensorimotor adaptation task.

The extent to which attention influences sensorimotor adaptation may also depend on the characteristics of the secondary task. Many secondary tasks, such as rapid serial visual presentation (18, 20), vocal shadowing (13), brightness discrimination, color discrimination, and auditory frequency discrimination (15, 19) require participants to detect subtle sensory signals within background noise. These tasks are considered data-limited because performance is predominantly restricted by the quality and discriminability of the sensory input, such as stimulus exposure duration, contrast, letter visibility, and masking conditions – rather than the amount of cognitive effort expended (35). When paired with a sensorimotor adaptation task, they are thought to draw on shared perceptual processing pathways essential for perceiving movement errors and discriminating sensory input relevant to the data-limited secondary task. Thus, the degree by which a secondary task influences sensorimotor adaptation likely depends on the extent of overlap in the sensory modalities and processing resources required for analyzing visual feedback, detecting movement errors, and executing corrective responses.

The effects of data-limited secondary tasks on sensorimotor adaptation have been mixed across studies, with evidence suggesting that interference is modulated by specific task characteristics. For example, auditory frequency discrimination has been shown to hinder sensorimotor adaptation during force field tasks (15) but does not appear to affect visuomotor adaptation (20, 21). In these cases, the secondary tasks differed in sensory modality and task nature (e.g., auditory discrimination vs. visuomotor adaptation). Theoretical accounts of cross-modal attention suggest that dividing attention between tasks that rely on the same sensory modality and task nature (i.e., cognitive operations), is more demanding than when dual-tasks are administered across sensory modalities and task natures, due to increased competition for overlapping perceptual and attentional resources (36). Thus, the degree to which a secondary task disrupts sensorimotor adaptation is likely to be most pronounced when the secondary task engages similar modality-specific and process-specific mechanisms as those required by the primary task.

To address the questions of whether and how dividing attention influences implicit sensorimotor adaptation, we employed a task paradigm typically referred to as error-clamped feedback, to emphasize implicit adaptation during visuomotor perturbation (32, 37). During the error-clamp perturbation, the angular trajectory of the feedback cursor is fixed along an invariant path, so that the cursor becomes rotated by a constant angle from the target location, irrespective of motor performance. As such, the cursor becomes directionally independent from the veridical reaching movement (32, 37–39). With perturbation onset, participants are made aware that the visual feedback is no longer under their control and are instructed to disregard it. This approach is intended to eliminate explicit re-aiming, thereby enabling an accurate assessment of implicit sensorimotor adaptation (32). For the present study, we coupled this approach with a visual random dot kinematogram (RDK) direction discrimination task (40), in order to measure how implicit sensorimotor adaptation is influenced by a secondary attention-demanding task. Unlike previously tested secondary tasks, that differed either in sensory modality or task nature from the sensorimotor adaptation task itself, both the RDK task and the implicit sensorimotor adaptation task engage similar visuospatial processes related to perceiving and planning movement direction in space. This provides an opportunity to examine whether interference in implicit motor adaptation arises from a shared reliance on perceptual mechanisms necessary for both error detection and the discrimination of task-relevant sensory signals. Further, we also present the attention-demanding secondary task during various stages of the movement itself (inward vs outward movement stages), in order to determine how the temporal specificity of the attentional distraction might influence error encoding during implicit sensorimotor adaptation.

## METHODS

### Participants

Forty-eight right-handed participants were recruited from the University of Queensland’s Department of Psychology and Macquarie University’s School of Psychological Sciences undergraduate participant pool (33 female, ages 17-32, mean age 21.6 ± 3). Handedness was determined using the Edinburgh Handedness Inventory (41). The number of participants per condition (n=12) was specified in advance based on previous studies with similar dual-task requirements and experimental design (17, 20, 21), and an estimated effect size of η2p > 0.26. These group sizes are also in accord with typical ranges used in other sensorimotor adaptation studies by various groups (1, 15, 16, 42). All students received course credit in exchange for their participation. The protocol was approved by the local ethics committee at The University of Queensland and Macquarie University. All participants provided written informed consent prior to participation.

### Apparatus

The experiment was conducted in a dimly lit room. Subjects sat at a custom tabletop workspace, where an LCD monitor (Dell P2417H) that displayed visual stimuli was placed horizontally to the floor. Using custom Matlab software, reaching behavior was recorded with a Wacom digitizing tablet and digitizing pen at 60 Hz (Intuos PTH-851/K1-CX). The digitizing tablet was placed underneath the monitor to occlude vision of the tablet and the participant’s arm, with the center of the tablet aligned with the center of the monitor. The center of the participant’s body was roughly aligned with the center of the digitizing tablet. A keyboard was placed beside the custom tabletop workspace, so participants could perform key-presses to indicate their response after the secondary attention-demanding task was presented. Participants held the digitizing pen with a power grip, and auditory tones were presented from a monitor speaker (Labtech LCS-1016).

### Random Dot Kinematogram Direction Discrimination Task

Participants performed a two-alternative forced-choice direction-discrimination task (40). During each trial, a center fixation (start) point (white empty circle - 1 cm in diameter) was presented in the center of the screen. Participants were instructed to maintain fixation on the centrally located circle, while continuously moving white dots (n=500) filled a large transparent ring with a diameter equal to the width of the computer screen (1080 pixels; 30 cm). The coherent motion stimulus was generated online using a standard random dot kinematogram (RDK) algorithm (40) and was presented on screen with a frame rate of 60 Hz. Based on pilot testing, hand movement duration for the ballistic movement task took approximately 300ms. Thus, on each trial, the RDK consisted of a 500ms sequence of video frames to ensure that participants had to engage in the discrimination task for the entire movement duration. During the motion presentation, each dot was sent to the video display for one video frame (16.67ms). To create apparent motion, a subset of the dots was displaced either upwards or downwards from their previous positions while the remaining dots were randomly repositioned. The participant’s task was to determine the direction of motion from the non-randomly positioned dots (traveling either up or down) and indicate their percept of directionality by pressing the corresponding arrow key on a keyboard with their non-dominant hand. This was prompted by displaying onscreen the text “Were the dots moving Up or Down?”, and the visual display froze until the participant made a response on the keyboard. For every RDK trial, auditory feedback was given, with a double beep indicating an incorrect response and a single beep signaling a correct response.

Prior to the sensorimotor adaptation task, 60 non-reaching trials were completed to determine an appropriate motion coherence (i.e., the percentage of coherently moving dots) for each participant in the dual-task conditions. We established the motion coherence at which participants could achieve a 60% accuracy in discrimination performance using the Bayesian adaptive staircasing method in Matlab’s QuestMean function (43), which is based on the QUEST algorithm (44). The motion coherence that best fit this accuracy estimate was then applied for the remainder of the experiment. Low motion coherence values, such as 0.01 coherently moving dots, represent an extremely difficult task, as only 1% of the dots are traveling in a consistent direction. Conversely, high motion coherence values, such as 0.90 coherently moving dots, represent an easier task, as 90% of the dots are traveling consistently in one direction. The 60% accuracy threshold was chosen to ensure that the discrimination task posed a significant level of difficulty to necessitate the engagement of substantial attentional resources; and it also aligns with the accuracy levels achieved by participants in previous studies (17–20).

### Visuomotor adaptation Task

Participants used a digitizing pen to control a circular white cursor (1 cm in diameter) displayed on the monitor with a default black background. To begin a trial, participants were instructed to place the cursor in the starting position, which was a white ring 1 cm in diameter. After holding the cursor in the starting position for 1 second, a green target circle 1 cm in diameter appeared at a distance of 7.5 cm in the 12 o’clock direction. Participants were then instructed to quickly reach with a center-out ballistic straight-line motion as if they were attempting to “slice through” the green target. Participants had 800ms to initiate and complete the movement; and were instructed to hold their hand stationary at the extent of their reach, until the feedback of their reach accuracy was no longer displayed. After completing the ballistic “outward movement” with the cursor displaying veridical online feedback, the endpoint location of the cursor froze on the display for 1 second, after which participants had 1 second to complete an “inward movement” returning to the starting position. Importantly, no online cursor feedback was provided during the inward movement stage. Instead, participants were guided back to the starting position by a white ring, whose radius represented the radial distance between the hand and the start point. Once the hand was within 0.5 cm of the start location, the ring transformed into cursor feedback. Participants were required to keep their hand in the starting position for 500ms before being prompted to "Press any key to continue." In dual-task trials, a different prompt appeared, asking: "Were the dots moving Up or Down?". Depending on the group (see General Experimental Procedures), the RDK could be displayed during either the *inward* or *outward* portion of the movement. To proceed, participants needed to press the corresponding up or down key. If the outward movement took longer than 800ms, or the inward movement took longer than 1 second, an error message of “Too slow” was presented.

To emphasize sensorimotor adaptation induced by implicit processes, we introduced error-clamped feedback (32) during the perturbation phase. Specifically, during the outward movement stage, the cursor extent remained yoked to the movement amplitude of the participant’s hand, while the cursor direction was fixed along an invariant path, 45° degrees away from the target location (counterbalanced - clockwise or counterclockwise across participants). For all experimental groups, the first trial of the perturbation was experienced by the participant, and then the nature of the perturbation was explained by the experimenter. Participants were informed that the cursor was now “clamped” to a fixed path that would miss the target location by 45°, no matter their actual reaching direction. Participants were then instructed to ignore the irrelevant feedback and continue reaching directly towards the target. To confirm the task instructions were fully understood, participants were required to explain to the experimenter how and where they should be aiming their reaches following first exposure. If the participant indicated they should aim anywhere but the target, the experimenter would then repeat the instruction until it was clearly understood.

### General Experimental Procedures

After establishing the motion coherence for each participant, participants performed the visuomotor adaptation task with or without the direction discrimination task (Figure 1), depending on the requirements for the different conditions and experiment phases (Figure 2). The aim of this study was twofold. First, we investigated how diverting attentional resources away from the primary visuomotor adaptation task influences implicit sensorimotor adaptation within an error clamped feedback paradigm. Second, we examined the temporal specificity of dividing attention during the encoding of error for implicit adaptation, by introducing the dual-task during varying portions of the movement (inward vs. outward). To achieve this, we conducted an experiment in which participants were randomly assigned to one of four groups in a 2x2 factorial design, with 12 participants per group: OutwardSingle-Dual (n=12); OutwardDual-Single (n=12); InwardSingle-Dual (n=12); and InwardDual-Single (n=12). This design involved two levels: the movement portion, in which the RDK dual-task was applied - ‘Outward’ and ‘Inward’ – and the order in which the participants performed the tasks - either beginning with the single implicit adaptation task, followed by the RDK dual-task (single-dual), or vice versa (dual-single).

**Figure 1.**
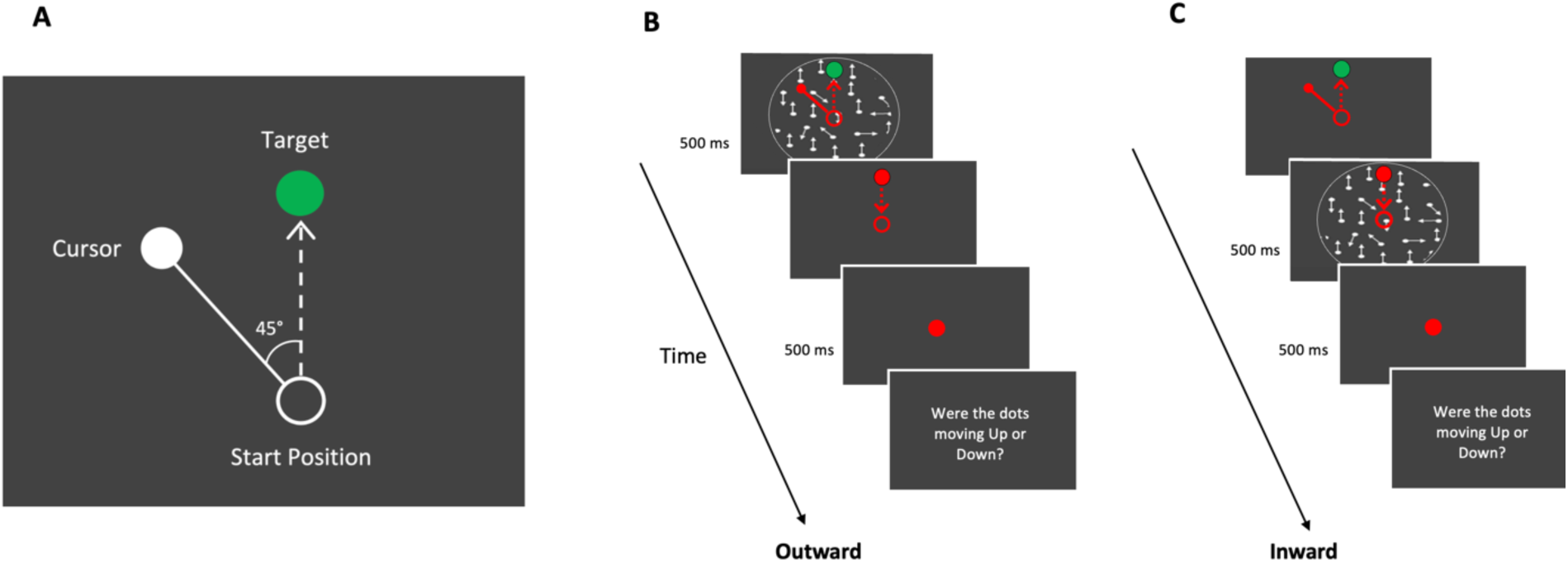
Schematic of the error-clamp paradigm and dual-task experimental design. (A) Participants performed center-out reaching movements from a start position (white circle) to a visual target (green circle) while controlling a white cursor. An error-clamp paradigm was used, in which the cursor was constrained to follow a fixed trajectory offset by 45° from the target direction (counterclockwise shown; direction was counterbalanced across participants). The dashed arrow indicates the instructed reach path, while the solid arrow represents the clamped cursor trajectory. (B) In the Outward dual-task condition, a random dot kinematogram (RDK) appeared during the outward reach, coinciding with the period of sensory prediction error associated with the clamped feedback. In the Inward dual-task condition, the RDK was instead presented during the return movement, after the error- clamp phase. Participants judged the global motion direction (upward or downward) of the dots in both dual-task conditions. In Single-task trials, the RDK was omitted and replaced with a neutral prompt (“Press any key to continue”). Red highlights in the figure are used solely for illustrative clarity; during the experiment, all cursor and movement elements were white.

**Figure 2.**
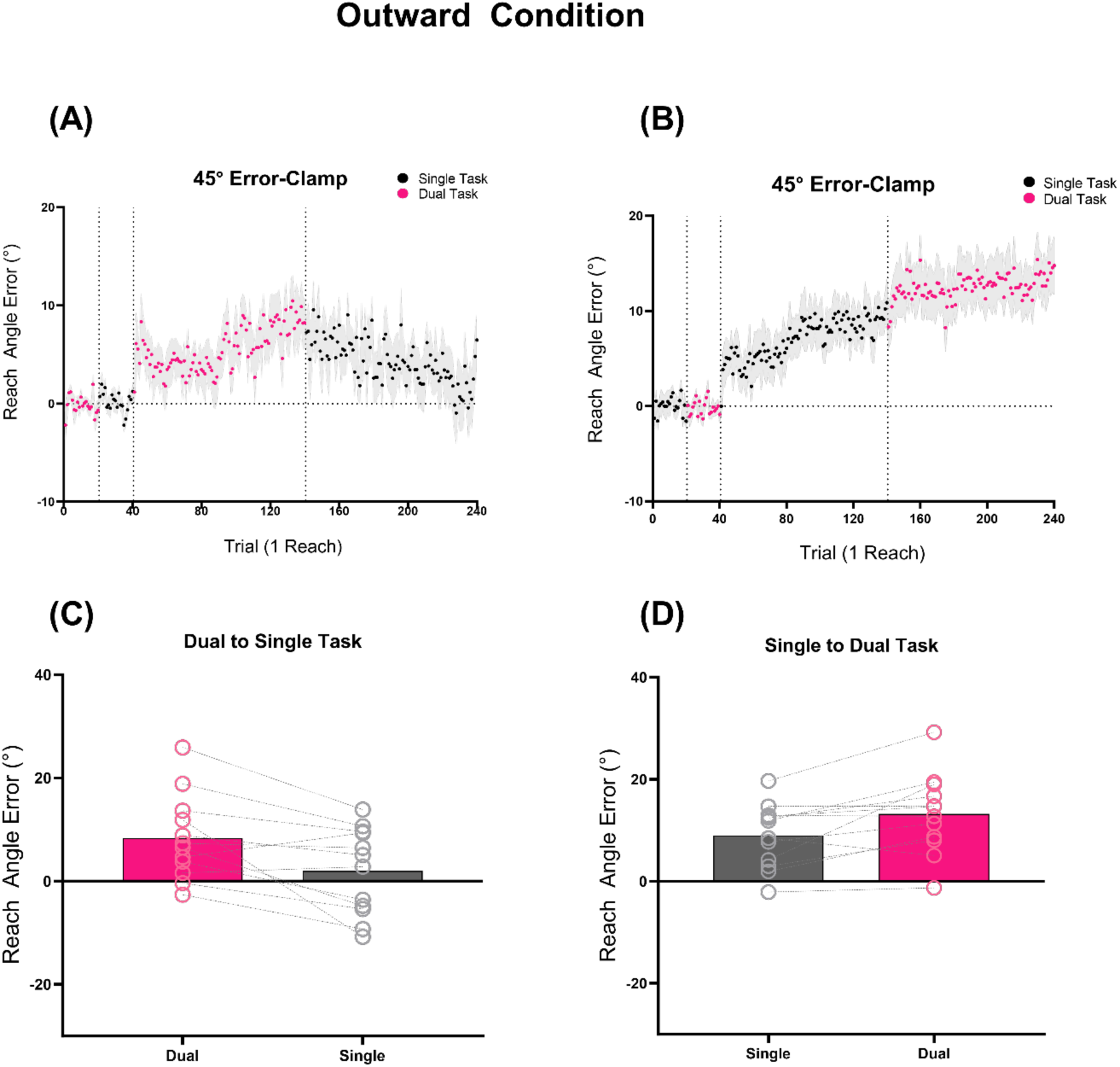
*Effects of dual-tasking on implicit adaptation during outward movement.* (A, B) Trial-by-trial reach angles during error-clamp trials for participants in the OutwardDual-Single (A) and OutwardSingle-Dual (B) groups. Participants experienced the RDK task during the outward phase of the reach in the dual-task blocks. Pink dots show reach angles during dual-task trials; black dots show reach angles during single-task trials (no RDK). Traces reflect group means (N = 12 per group), with shaded areas representing ±1 SEM. Vertical dotted lines indicate transitions between baseline, dual-task, and single-task blocks. (C, D) Mean reach angle over the final 20 error-clamp trials in each task phase for the OutwardDual-Single (C) and OutwardSingle-Dual (D) groups. Each circle represents a participant’s mean reach angle. Horizontal bars indicate group means; error bars show ±1 SEM. Grey lines connect individual participants across conditions, illustrating within-subject differences.

To start the experiment, all participants underwent a familiarization phase, where they first performed 10 dual-task reaches to the target, followed by 10 single-task reaches to the target (with order counterbalanced across groups). Next, in a baseline phase, participants completed 20 dual-task baseline reaches, followed by 20 single-task baseline reaches. All these trials were completed with veridical online cursor feedback. Subsequently, for the first perturbation phase, participants in the OutwardDual-Single and InwardDual-Single groups completed 100 trials during the dual-task block. Here, the error-clamp perturbation was paired with the RDK dual-task during the outward or inward movements, respectively (also counterbalanced by group). Next, during the second perturbation phase, participants transitioned into the single-task block, where they performed 100 trials with only the error-clamp perturbation applied. Conversely, participants in the OutwardSingle-Dual and InwardSingle-Dual groups completed the tasks in the reverse sequence, always performing the single-task first, followed by the dual-task.

### Data Analysis

Hand kinematics and direction discrimination task responses were analyzed offline in Matlab. For all trials, the angular difference between the target direction and the direction of the vector from the participants’ starting position to the point where their movements passed a radial distance of 7.5 cm, was calculated as the terminal reach-angle. Terminal reach angles from participants experiencing clockwise perturbations were reversed so that positive angles uniformly reflect gains in adaptation magnitude. Reach angles that were more than 3 standard deviations away from the mean during each phase of the experiment were removed as outliers (total proportion of trials removed = 0.18%). Reaction time was measured as the duration between the target’s appearance and the point at which the cursor’s position exceeded 5% of the required target distance. Movement duration was defined as the time taken for the cursor to travel from 5% to 100% of the target distance. Baseline performances were quantified by averaging the terminal reach-angles for the 20 baseline trials in the single-task and dual-task blocks respectively. For the perturbation phase, terminal reach-angles for the final 20 trials were averaged in the single-task and dual-task blocks respectively, to quantify the degree of implicit adaptation.

### Statistical Analysis

To investigate the degree of implicit sensorimotor adaptation under different attentional requirements, we analyzed the average terminal reach-angle from the baseline and perturbation phases for all groups (OutwardDual-Single; OutwardSingle-Dual; InwardDual-Single and InwardSingle-Dual). We subjected mean terminal reach-angles from the final 20 trials of the perturbation phase, as well as the 20 baseline trials for both the single-task and dual-task blocks, to separate mixed factorial 3-way ANOVAs for the Inward and Outward groups. Specifically, the Task Order (Dual-Single task order, Single-Dual task order) was used as a between-subjects factor, while Task Nature (Dual Task, Single Task) and Task Phase (Baseline phase, Perturbation phase) were used as within-subjects factors. To directly assess how dividing attention influenced sensorimotor adaptation, we also conducted planned comparisons within each group. Specifically, we tested whether the amount of implicit sensorimotor adaptation changed when participants transitioned from dual-task to single-task conditions (i.e., first vs second perturbation phase) or vice versa. All figures are presented as mean ± standard error of mean. We also calculated effect sizes using the partial eta square (η²p). All statistical analyses were performed in jamovi (version 2.5)

## RESULTS

### RDK Accuracy

Participants’ ability to discern the direction of random-dot motion from varying motion-strengths was assessed before the experiment. The motion coherence that resulted in 60% correct discrimination was selected for the dual-task blocks of the main experiment. The average motion coherence was 0.067±0.017, 0.082±0.016, 0.06±0.01 and 0.064±0.01 for the OutwardDual-Single,OutwardSingle-Dual, InwardDual-Single and InwardSingle-Dual groups respectively. In the reaching experiment, participants performed the direction discrimination task concurrently with the sensorimotor adaptation task for the baseline and perturbation phase dual-task blocks. To determine whether RDK task performance accuracy varied between the baseline and perturbation phases, we conducted a mixed factorial ANOVA, treating phase (baseline vs. perturbation) as a within-subject factor and movement stage (outward vs. inward) as a between-subject factor. We found that RDK accuracy for the baseline phase (averaged across the 20 baseline trials) did not differ significantly from that of the perturbation phase (averaged across the 100 training trials; no main effect of phases: f(1,44) = 0.0376, p = 0.847, η²p = 0.001). In addition, there were no differences in RDK accuracy irrespective of whether the RDK was presented during the outward or the inward portion of the movement for the Outward and Inward groups respectively (no main effect of groups: f(3,44) = 0.506, p = 0.68, η²p = 0.033). On average, the RDK accuracies for OutwardDual-Single, OutwardSingle-Dual, InwardDual-Single and InwardSingle-Dual groups were 63.5±3%, 68.5±4%, 64.1±3% and 67.4±3% (chance level = 50%) respectively. These accuracy levels are consistent with those observed in prior dual-task studies involving data-limited tasks such as rapid serial visual presentation, brightness discrimination, and sound discrimination tasks (17–21). This indicates that participants diverted substantial attention to the direction discrimination task, thereby reducing the attentional resources available for simultaneous implicit sensorimotor adaptation.

### Effects of Attentional Distraction during Outward Movement on Implicit Adaptation

We first investigated whether implicit sensorimotor adaptation is influenced by a secondary attention-dividing task that engages overlapping visuospatial processes. To achieve this, we presented a visual random dot kinematogram (RDK) direction discrimination task together with the sensorimotor adaptation task during the outward portion of the movement (movements toward the target). Unlike many prior dual-task studies that used secondary tasks differing in sensory modality or cognitive demands, both the RDK and sensorimotor adaptation tasks in the present study required processing of movement direction in visual space. This allowed us to test whether competition for shared perceptual mechanisms — specifically those involved in detecting movement error and visual motion discrimination — would influence implicit sensorimotor adaptation. We compared two groups of participants: one that experienced a 45° error-clamp under dual-task conditions before switching to single-task conditions (OutwardDual-Single group) and another that experienced the reverse order (OutwardSingle-Dual group). If the RDK task disrupts movement error processing, we expected reduced sensorimotor adaptation magnitudes during dual-task conditions, and greater adaptation when the secondary task is removed and attention is no longer divided.

During the perturbation phase, the introduction of error-clamped feedback resulted in terminal reach-angle changes to compensate for the visual discrepancy for both the OutwardDual-Single (Figure 2A) and OutwardSingle-Dual (Figure 2B) groups. This was confirmed statistically by the main effect of Phase (f(1,22) = 32.6, p < 0.001, η²p = 0.597), implying that the introduction of error-clamped feedback resulted in implicit sensorimotor adaptation. Importantly, the degree of implicit sensorimotor adaptation was strongly influenced by whether attention was divided during the perturbation phase (main effect of task nature: f(1,22) = 14.7, p < 0.001, η²p = 0.401 and phase*task nature interaction: f(1,22) = 20.9, p < 0.001, η²p = 0.487). Interestingly, contrary to our initial expectations, we observed that implicit sensorimotor adaptation was more pronounced when attention was divided under dual-task conditions for both groups (OutwardDual-Single: 8.4±2.4°, OutwardSingle-Dual : 13.2±2.4°); in comparison, implicit sensorimotor adaptation was lower for single-task conditions, when attention was not divided (OutwardDual-Single: 2±2.4°, OutwardSingle-Dual: 8.9±1.8°).

To directly test the effect of changing attentional demands on implicit sensorimotor adaptation within individuals, we conducted planned contrasts comparing sensorimotor adaptation across dual-task and single-task phases separately for both the OutwardDual-Single and OutwardSingle-Dual groups. In the OutwardDual-Single group, the planned contrast revealed a reduction in implicit adaptation when participants transitioned from dual-task to single-task conditions, with adaptation levels approaching baseline by the end of the perturbation phase (t(11) = 3.24, p = 0.004, Figure 2C). Conversely, in the OutwardSingle-Dual group, the planned contrast indicated a significant increase in adaptation when participants transitioned from single-task to dual-task conditions, reaching adaptation levels significantly greater than in the single-task (t(11) = 3.38, p = 0.003, Figure 2D). This suggests that changes in attentional context—specifically, the addition or removal of attentional demands—directly modulate the magnitude of implicit sensorimotor adaptation. In particular, the addition of an attention dividing secondary task facilitated implicit sensorimotor adaptation relative to the level of adaptation observed during single-task performance. This finding represents a surprising departure from previous visuomotor adaptation studies, which report that dividing attention with a dual-task either has no effect on sensorimotor adaptation, or is disruptive when transitioning into a different attentional context (17-21). Here, we observed facilitation when attention was divided, highlighting that dividing attention can have differing effects on implicit sensorimotor adaptation.

### Effects of Attentional Distraction during Inward Movement on Implicit Adaptation

To determine whether the facilitative effect of dual-tasking depends on its timing relative to movement error processing, we presented the RDK task during the returning (inward) portion of the movement, following the outward reach and error exposure. If the temporal overlap between dividing attention and movement error encoding is critical, we would not expect to see any differences in the pattern of implicit adaptation regardless of the attentional context. On the other hand, if the timing of divided attention and movement error encoding is not essential, we would anticipate a similar pattern of results for dual and single-task conditions.

Figure 3A and 3B illustrates the pattern of implicit sensorimotor adaptation for both the InwardDual- Single and InwardSingle-Dual groups. Both InwardDual-Single and InwardSingle-Dual groups adapted their terminal reach-angles in response to the error-clamped feedback (main effect of phase: f(1,22) = 11.3, p = 0.003, η²p = 0.34). Interestingly, there was no evidence that sensorimotor adaptation under dual-task conditions had any effect on adaptation magnitude when the secondary discrimination task was performed during the inward (return to home) portion of the movement (no main effects of Task Nature: f(1,22) = 0.004, p = 0.946, η²p = 0 and Task Order f(1,22) = 0.98, p = 0.332, η²p = 0.043). We observed that the amount of sensorimotor adaptation did not significantly differ during the last 20 perturbation trials for the dual-task blocks (InwardDual-Single: 10.1±2.5°, InwardSingle-Dual: 5.99±3.7°) or the single-task blocks (InwardDual-Single: 11.4±2.8°, InwardSingle-Dual : 4.77±4.9°) for both groups. We also observed no task-nature*phase*group interaction effect (f(1,22) = 1.1, p = 0.314, η²p = 0.046). Thus, introducing the RDK task during the inward portion of the movement had little effect on sensorimotor adaptation, regardless of whether participants transitioned from dual-task to single-task conditions or the reverse. This highlights that the facilitatory effect of dual-tasking on implicit sensorimotor adaptation depends critically on its temporal alignment with the period of movement error processing.

**Figure 3.**
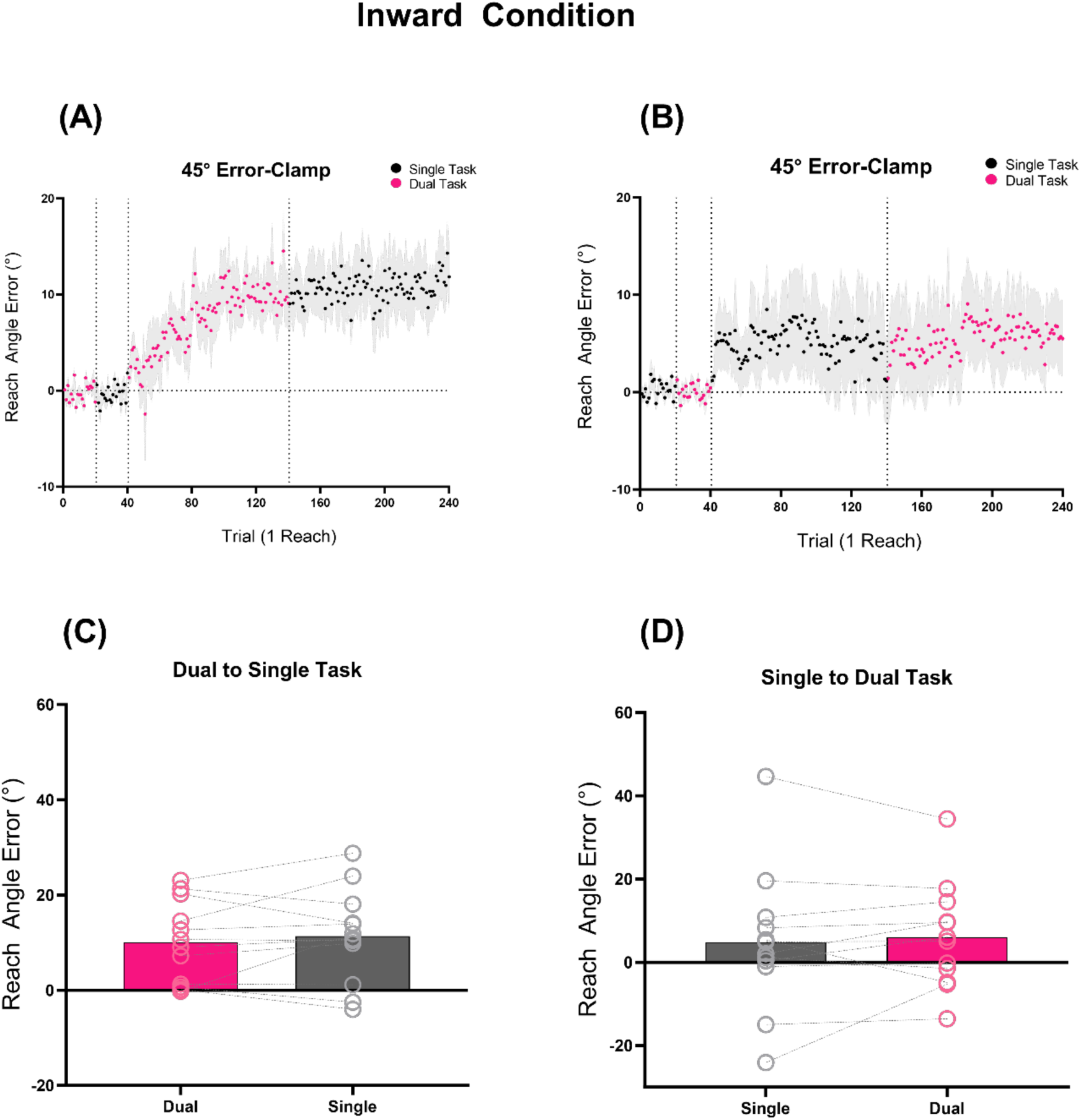
*Effects of dual-tasking on implicit adaptation during inward movement.* (A, B) Trial-by-trial reach angles during error-clamp trials for participants in the InwardDual-Single (A) and InwardSingle-Dual (B) groups. Participants experienced the RDK task during the inward phase of the reach in the dual-task blocks. Pink dots show reach angles during dual-task trials; black dots show reach angles during single-task trials (no RDK). Traces reflect group means (N = 12 per group), with shaded areas representing ±1 SEM. Vertical dotted lines indicate transitions between baseline, dual-task, and single-task blocks. (C, D) Mean reach angle over the final 20 error-clamp trials in each task phase for the InwardDual-Single (C) and InwardSingle-Dual (D) groups. Each circle represents a participant’s mean reach angle. Bars indicate group means; error bars show ±1 SEM. Grey lines connect individual participants across conditions, illustrating within-subject differences.

### Performing the RDK task did not interfere with reaction or movement times

To evaluate whether the presence of a secondary task altered basic temporal features of the reaching movement—potentially reflecting a shift in task approach that could account for differences in adaptation—we examined participants’ reaction times (RT) and movement times (MT) during the perturbation phase. Specifically, we asked whether performing the dual-task affected how participants initiated or executed their movements. This might imply the adoption of different strategies or altered attentional allocation. We performed a mixed factorial ANOVA with Task Nature (Single, Dual) as the within-subject factor, and Groups (Inward, Outward) and Task Order (Dual-Single, Single-Dual) as between subject factors.

We found no differences in RT across the groups for the single-task or dual-task blocks during the perturbation phase. The ANOVA revealed no main effect of Task Nature (f(1,44) = 0.173, p = 0.679, η²p = 0.004), Groups (f(1,44) = 0.61, p = 0.44, η²p = 0.014), or Task Order (f(1,44) = 1.81, p = 0.185, η²p = 0.04), and no task-nature*movement stage*task order interaction (f(1,44) = 1.754, p = 0.192, η²p = 0.038) .

A similar pattern of results was observed for movement times. There were no differences in MTs across the OutwardSingle-Dual; OutwardDual-Single; InwardSingle-Dual; and InwardDual-Single groups. The ANOVA revealed no main effect of Task Nature (f(1,44) = 0.154, p = 0.697, η²p = 0.003), Groups (f(1,44) = 0.0003, p = 0.986, η²p = 0.00), or Task Order (f(1,44) = 1.63, p = 0.208, η²p = 0.04), and no task-nature*groups*task order interaction (f(1,44) = 0.702, p = 0.406, η²p = 0.016). These results suggest that all groups used similar movement kinematics irrespective of whether they experienced the RDK task or not (single-task vs dual-task), as well as *when* they performed the RDK task (either inward or outward).

These results indicate that participants maintained consistent reaction and movement timing regardless of whether they were performing a dual-task or single-task, and regardless of when the RDK task was introduced (outward vs. inward movement). Together, these findings suggest that the differences in implicit sensorimotor adaptation observed in earlier analyses were not driven by changes in movement execution strategies, but rather by how changing attentional demands interacted with movement error processing.

## DISCUSSION

The current study was designed to examine whether and how attention influences implicit sensorimotor adaptation. Prior research yielded conflicting results. Some studies reported that dividing attention using a secondary task impaired sensorimotor adaptation (9–11, 15, 16), whereas other studies found no impairment (17–21). Because sensorimotor adaptation is a multifaceted process, we propose that these discrepant results may stem from differential influences of attention on the explicit and implicit components involved in sensorimotor adaptation tasks. Crucially, previous dual-task studies did not distinguish between explicit and implicit processes during sensorimotor adaptation, thus making it unclear whether the data reflect the influence of explicit re-aiming or implicit sensorimotor adaptation. Here we assessed the effect of attention on the implicit component of sensorimotor adaptation using the error-clamp paradigm (32). Surprisingly, we found a paradoxical enhancement of adaptation when participants simultaneously performed an attention-demanding motion discrimination task. This effect was robust and clear across most individuals relative to their single-task performance, irrespective of whether they performed the single-task first (OutwardSingle-Dual) or transitioned from the dual-task back to the single-task condition (OutwardDual-Single). Importantly, implicit sensorimotor adaptation was only enhanced when the attention task was executed during outward movements; while implicit sensorimotor adaptation remained unchanged when the attention task was performed as participants returned their hand to the starting position between reaches. This implies that the timing of attentional demands relative to movement error encoding may play a crucial role in influencing how attention affects the magnitude of implicit sensorimotor adaptation.

### Task-Specific Resource Limitations in Dual-Task Paradigms

Previous studies that employ data-limited secondary tasks, such as rapid serial visual presentation (RSVP), vocal shadowing, and visual or auditory discrimination tasks (13, 15, 18–20), have typically interpreted attentional effects on sensorimotor adaptation through the lens of fixed-capacity resource models (45). According to fixed capacity resource models, attentional resources are limited and cannot be simultaneously allocated to multiple demanding tasks without incurring performance costs. Consequently, the extent to which the secondary task influences sensorimotor adaptation depends on the competition for limited-capacity attentional resources between the primary and secondary tasks. For the Outward condition, we observed that implicit sensorimotor adaptation was greater when individuals simultaneously performed both the RDK and sensorimotor adaptation tasks, relative to when they performed the sensorimotor task in isolation. Importantly, this occurred without deterioration to the secondary motion discrimination task, suggesting that dual-task performance is not necessarily restricted by shared capacity limits induced by visuospatial perceptual demands. If such a constraint had played a major role in this context, then either RDK accuracy or the magnitude of implicit sensorimotor adaptation should have deteriorated during these dual-task trials.

### Mechanisms underlying attentional modulation of implicit sensorimotor adaptation

The present findings demonstrate that divided attention can influence implicit sensorimotor adaptation in ways that are not easily accounted for by conventional shared capacity resource models. This prompts the critical question of what underlying mechanisms support this interaction? The fact that participants exhibited greater implicit sensorimotor adaptation during the dual-task trials compared to the single-task trials for the Outward but not the Inward condition suggests a modulation of the ability to detect sensory prediction errors and/or transform them into appropriate adaptive responses. Therefore, this effect could in principle be mediated by both “low-level” sensorimotor factors, and “high-level” cognitive mechanisms. A low-level explanation is attractive, as both the RDK task and the implicit sensorimotor adaptation task engage similar visuospatial processes related to perceiving and planning motion direction in space. This stands in contrast to prior studies utilizing other motor adaptation dual-tasks —such as RSVP (18, 20), vocal shadowing (13), and tasks involving brightness, color, and auditory frequency discrimination (15, 19) — which generally observed either impairments (15) or no effect on sensorimotor adaptation (13, 18–20).

One potential low-level explanation is that the visuospatial characteristics of the RDK task might have modulated the salience of the cursor by increasing its perceptual contrast against the dynamic background. This type of salience boost would be unlikely during inward movements, as the RDK was presented only after the error-clamped cursor had disappeared. This interpretation aligns with the idea that the nervous system actively estimates the relevant source of motor errors to guide adaptation (46). In support of this, previous works (47, 48) have shown that when participants were specifically instructed to attend to, or ignore, particular cursors during a multi- cursor paradigm, implicit sensorimotor adaptation occurred only for the relevant cursor, while no adaptation was observed for the irrelevant ones.

Another possible explanation for instances of dual-task enhanced implicit adaptation relates to higher-level cognitive processes, such as the selective allocation of attentional resources (36). For instance, when performing the RDK and sensorimotor adaptation tasks concurrently, attention may be oriented more toward the RDK task rather than the sensorimotor adaptation task. This shift in attentional focus could engage processes that enhance implicit sensorimotor adaptation by reducing conscious interference, promoting reliance on automatic and implicit mechanisms. Notably, these mechanistic accounts may help to reconcile the discrepant findings reported by previous studies that examine the effects of dual-task interference on sensorimotor adaptation.

Comparison with previous work shows that the increased attentional demands of a dual-task either impairs or does not affect sensorimotor adaptation (15–21). This suggests that if a differential engagement of cognitive strategies contributes to our observed dual-task benefit, such effects may require a close alignment in attentional demands between the primary and secondary tasks. To further clarify this finding, future studies should directly compare secondary tasks involving visuospatial processes related to perceiving and planning motion direction in space (as in the current study), with secondary tasks containing a less specific relationship to the reaching task (i.e., salient and challenging non-visual stimuli). It will also be important to manipulate the perceptual difficulty of the RDK task in order to systematically vary attentional load. If increased attentional engagement facilitates implicit sensorimotor adaptation, lower-coherence (i.e., more demanding) stimuli should enhance sensorimotor adaptation, whereas higher-coherence (i.e., less demanding) stimuli should attenuate this effect.

Arousal is another high-level factor that may influence the dual-task effects observed in the present study. In particular, the performance of a cognitively demanding task (such as dual-tasking) has been associated with increased autonomic arousal measures (such as a larger pupil size, decreased pulse amplitude, and skin conductance) when compared to a less demanding task (53). Such fluctuations in autonomic measures are established psychophysiological markers of cognitive load and arousal (54, 55). In our task, increased attentional demands under the dual-task condition may have modulated arousal in a way that enhanced the encoding of sensory prediction errors for the OutwardSingle-Dual group, ultimately leading to increased implicit adaptation. This interpretation is partly informed by findings from Tomassi et al. (56) who showed that increased autonomic arousal was associated with greater adaptation in a vocal-auditory-motor adaptation task. However, as that study did not distinguish between explicit and implicit components of adaptation, it remains uncertain whether the observed learning enhancement was specifically attributable to arousal during implicit sensorimotor adaptation (in particular, the processing of sensory prediction errors). Given that a dual-task of sufficient task-difficulty should increase arousal, irrespective of the precise nature of the secondary task, the fact that previous dual-task sensorimotor adaptation studies not distinguishing between implicit and explicit adaptation processes show either impairment or no effect on adaptation, implies that an effect of arousal on sensorimotor adaptation should relate to implicit adaptation. This hypothesis could be directly evaluated in future work by incorporating pupillometric or other autonomic measures within paradigms that emphasize implicit adaptation, such as the error-clamp method employed here.

### Choking in High Pressure Situations and External Focus of Attention during Skill Acquisition

In daily activities, individuals often need to adjust and refine movements in response to changing environmental conditions or internal states. This typically occurs while we face distractions from various sources that divert attention away from the main task. Such distractions are often viewed as obstacles to effective sensorimotor adaptation. However, our findings suggest that these distractions are not always detrimental. Instead, we observe a counterintuitive effect: diverting attention with a secondary task can facilitate implicit sensorimotor adaptation, relative to an equal sized perturbation encoded without attentional distraction.

A similar paradox has been noted in the sports science literature in relation to choking under high-pressure situations. The phenomenon of “choking” refers to ‘an acute and considerable decrease in skill execution and performance when self-expected standards are normally achievable, which is the result of increased anxiety under perceived pressure’ (57). An example of this could be a top NBA player who usually makes 95% of their free throws, but then misses two critical free throws in the Finals that could have tied the game. One explanation for this phenomenon is that heightened pressure induces an internal focus of attention that disrupts the automaticity of well-practiced skills (58). Skilled movements are typically executed with minimal conscious control. However, because the stakes are so high, individuals may turn their attention to the execution of their movements and attempt to consciously regulate them, which ironically leads to worse performance. Supporting this idea, there is evidence that experienced athletes tend to perform better when their attention is shifted toward an external event rather than directing attention onto their own movements (59). Relating to our results, this would allow for explicit attention processes to reduce their interference with well refined, experience dependent sensorimotor sequences (as depicted in Figure 2B).

Interestingly, these shifts in attentional focus not only affect performance, but also play a crucial role in the learning process. Research on skill acquisition has shown that the way learners allocate their attention can significantly influence how they acquire and refine skills over time (60). Specifically, directing attention from the movements themselves to the environment—has been found to enhance learning (61). In a series of studies involving different movement skills, like skiing, golf pitch, and tennis backhand, Wulf and colleagues found that instructions encouraging an external focus of attention, which directs learners’ attention to the outcome of their actions, results in more effective learning compared to instructions that focus on the movements themselves (62–64). These results suggest that the way attention is directed – whether towards the execution of movements, or the effects of movements on the environment – plays a pivotal role in motor performance and skill acquisition. Taken together, this aligns with the central finding of our study, which suggests that allocating attentional resources to a motor task does not always enhance performance, adaptation, or learning. In some cases, increased attentional resources can hinder these processes.

## Conclusion

In conclusion, the present work provides new insight into the complex relationship between attention and implicit sensorimotor adaptation. Our results show that people exhibit greater implicit sensorimotor adaptation when their attention is divided by the concurrent execution of a secondary visuospatial task. However, this paradoxical effect was only present when attention was divided during the Outward movement stage and not the Inward movement stage. This suggests that the timing of divided attention relative to movement error encoding may play a crucial role in influencing how attention affects the magnitude of implicit sensorimotor adaptation. In general terms, these results are difficult to reconcile with the notion that implicit motor adaptation is predominantly automatic and requires minimal attention, because attentional modulation clearly influences the extent of sensorimotor adaptation. In practical terms, these findings highlight the potential for enhancing sensorimotor adaptation, as well as the performance and learning of various movement skills, by strategically managing attentional focus.

## Acknowledgements

TJC is funded by the Australian Research Council (DP230102179). EP is funded by Macquarie University Performance and Expertise Centre Seeding Grant.

## References

1. Krakauer JW, Pine ZM, Ghilardi MF, Ghez C. Learning of visuomotor transformations for vectorial planning of reaching trajectories. Journal of Neuroscience. 2000;20(23):8916–24.

2. Shadmehr R, Mussaivaldi FA. Adaptive Representation of Dynamics during Learning of a Motor Task. Journal of Neuroscience. 1994;14(5):3208–24.

3. McGuire LM, Sabes PN. Sensory transformations and the use of multiple reference frames for reach planning. Nat Neurosci. 2009;12(8):1056–61.

4. Sabes PN. The planning and control of reaching movements. Curr Opin Neurobiol. 2000;10(6):740–6.

5. Sober SJ, Sabes PN. Multisensory integration during motor planning. J Neurosci. 2003;23(18):6982–92.

6. Sober SJ, Sabes PN. Flexible strategies for sensory integration during motor planning. Nat Neurosci. 2005;8(4):490–7.

7. Curran T, Keele SW. Attentional and Nonattentional Forms of Sequence Learning. J Exp Psychol Learn. 1993;19(1):189–202.

8. Nissen MJ, Bullemer P. Attentional Requirements of Learning - Evidence from Performance-Measures. Cognitive Psychol. 1987;19(1):1–32.

9. Redding GM, Clark SE, Wallace B. Attention and prism adaptation. Cogn Psychol. 1985;17(1):1–25.

10. Redding GM, Rader SD, Lucas DR. Cognitive Load and Prism Adaptation. J Mot Behav. 1992;24(3):238–46.

11. Redding GM, Wallace B. Cognitive interference in prism adaptation. Percept Psychophys. 1985;37(3):225–30.

12. Eversheim U, Bock O. Evidence for processing stages in skill acquisition: a dual-task study. Learn Mem. 2001;8(4):183–9.

13. Galea JM, Sami SA, Albert NB, Miall RC. Secondary tasks impair adaptation to step- and gradual-visual displacements. Exp Brain Res. 2010;202(2):473–84.

14. Ingram HA, van Donkelaar P, Cole J, Vercher JL, Gauthier GM, Miall RC. The role of proprioception and attention in a visuomotor adaptation task. Exp Brain Res. 2000;132(1):114–26.

15. Taylor JA, Thoroughman KA. Divided attention impairs human motor adaptation but not feedback control. Journal of Neurophysiology. 2007;98(1):317–26.

16. Taylor JA, Thoroughman KA. Motor Adaptation Scaled by the Difficulty of a Secondary Cognitive Task. Plos One. 2008;3(6).

17. Bedard P, Song JH. Attention modulates generalization of visuomotor adaptation. J Vision. 2013;13(12).

18. Im HY, Bedard P, Song JH. Encoding attentional states during visuomotor adaptation. J Vision. 2015;15(8).

19. Im HY, Bedard P, Song JH. Long Lasting Attentional-Context Dependent Visuomotor Memory. J Exp Psychol Human. 2016;42(9):1269–74.

20. Song JH, Bedard P. Paradoxical Benefits of Dual-Task Contexts for Visuomotor Memory. Psychol Sci. 2015;26(2):148–58.

21. Wang TSL, Song JH. Impaired visuomotor generalization by inconsistent attentional contexts. Journal of Neurophysiology. 2017;118(3):1709–19.

22. Taylor JA, Ivry RB. The role of strategies in motor learning. Ann N Y Acad Sci. 2012;1251:1–12.

23. Taylor JA, Krakauer JW, Ivry RB. Explicit and implicit contributions to learning in a sensorimotor adaptation task. J Neurosci. 2014;34(8):3023–32.

24. McDougle SD, Bond KM, Taylor JA. Explicit and Implicit Processes Constitute the Fast and Slow Processes of Sensorimotor Learning. J Neurosci. 2015;35(26):9568–79.

25. Schween R, McDougle SD, Hegele M, Taylor JA. Assessing explicit strategies in force field adaptation. J Neurophysiol. 2020;123(4):1552–65.

26. Redding GM, Wallace B. Adaptive spatial alignment and strategic perceptual-motor control. J Exp Psychol Hum Percept Perform. 1996;22(2):379–94.

27. Mazzoni P, Krakauer JW. An implicit plan overrides an explicit strategy during visuomotor adaptation. J Neurosci. 2006;26(14):3642–5.

28. Tseng YW, Diedrichsen J, Krakauer JW, Shadmehr R, Bastian AJ. Sensory prediction errors drive cerebellum-dependent adaptation of reaching. J Neurophysiol. 2007;98(1):54–62.

29. Bae GY, Luck SJ. Perception of opposite-direction motion in random dot kinematograms. Vis cogn. 2022;30(4):289–303.

30. Langsdorf L, Goehringer F, Schween R, Schenk T, Hegele M. Additional cognitive load decreases performance but not adaptation to a visuomotor transformation. Acta Psychol (Amst). 2022;226:103586.

31. Kagerer FA, Contreras-Vidal JL, Stelmach GE. Adaptation to gradual as compared with sudden visuo-motor distortions. Exp Brain Res. 1997;115(3):557–61.

32. Morehead JR, Taylor JA, Parvin DE, Ivry RB. Characteristics of Implicit Sensorimotor Adaptation Revealed by Task-irrelevant Clamped Feedback. J Cognitive Neurosci. 2017;29(6):1061–74.

33. Bond KM, Taylor JA. Flexible explicit but rigid implicit learning in a visuomotor adaptation task. J Neurophysiol. 2015;113(10):3836–49.

34. Stadler M. Role of Attention in Implicit Learning. J Exp Psychol: Learning, Memory and Cognition. 1995;21(3):674–85.

35. Norman DA, Bobrow DG. On data-limited and resource-limited processes. Cognitive Psychol. 1975;7(1):44–64.

36. Treisman AM, Davies A. Divided Attention to Ear and Eye. In: Wolfe J, Robertson L, editors. From Perception to Consciousness: Searching with Anne Treisman: Oxford University Press; 2012. p. 0.

37. Kim HE, Parvin DE, Ivry RB. The influence of task outcome on implicit motor learning. Elife. 2019;8.

38. Vandevoorde K, Orban de Xivry JJ. Internal model recalibration does not deteriorate with age while motor adaptation does. Neurobiol Aging. 2019;80:138–53.

39. Vaswani PA, Shmuelof L, Haith AM, Delnicki RJ, Huang VS, Mazzoni P, et al. Persistent residual errors in motor adaptation tasks: reversion to baseline and exploratory escape. J Neurosci. 2015;35(17):6969–77.

40. Roitman JD, Shadlen MN. Response of neurons in the lateral intraparietal area during a combined visual discrimination reaction time task. J Neurosci. 2002;22(21):9475–89.

41. Oldfield RC. The assessment and analysis of handedness: the Edinburgh inventory. Neuropsychologia. 1971;9(1):97–113.

42. Wu HG, Smith MA. The generalization of visuomotor learning to untrained movements and movement sequences based on movement vector and goal location remapping. J Neurosci. 2013;33(26):10772–89.

43. King-Smith PE, Grigsby SS, Vingrys AJ, Benes SC, Supowit A. Efficient and unbiased modifications of the QUEST threshold method: theory, simulations, experimental evaluation and practical implementation. Vision Res. 1994;34(7):885–912.

44. Watson AB, Pelli DG. QUEST: a Bayesian adaptive psychometric method. Percept Psychophys. 1983;33(2):113–20.

45. Kahneman D. Attention and effort. prentice-Hall; 1973.

46. Berniker M, Kording K. Estimating the sources of motor errors for adaptation and generalization. Nat Neurosci. 2008;11(12):1454–61.

47. Kasuga S, Hirashima M, Nozaki D. Simultaneous processing of information on multiple errors in visuomotor learning. Plos One. 2013;8(8):e72741.

48. Tsay J, Parvin DE, Dang KV, Stover AR, Ivry RB, Morehead JR. Implicit Adaptation Is Modulated by the Relevance of Feedback. J Cogn Neurosci. 2024;36(6):1206–20.

49. Poh E, Al-Fawakhiri N, Tam R, Taylor JA, McDougle SD. Top-down effects in motor generalization. bioRxiv. 2022.

50. McDougle SD, Taylor JA. Dissociable cognitive strategies for sensorimotor learning. Nat Commun. 2019;10.

51. Leow LA, Gunn R, Marinovic W, Carroll TJ. Estimating the implicit component of visuomotor rotation learning by constraining movement preparation time. J Neurophysiol. 2017;118(2):666–76.

52. Haith AM, Huberdeau DM, Krakauer JW. The influence of movement preparation time on the expression of visuomotor learning and savings. J Neurosci. 2015;35(13):5109–17.

53. Tursky B, Shapiro D, Crider A, Kahneman D. Pupillary, heart rate, and skin resistance changes during a mental task. J Exp Psychol. 1969;79(1):164–7.

54. Kahneman D, Beatty J. Pupil diameter and load on memory. Science. 1966;154(3756):1583-5.

55. Reimer J, McGinley MJ, Liu Y, Rodenkirch C, Wang Q, McCormick DA, et al. Pupil fluctuations track rapid changes in adrenergic and cholinergic activity in cortex. Nat Commun. 2016;7:13289.

56. Tomassi NE, Turashvili DM, Williams A, Walsh B, Stephen EP, Stepp CE. Investigating Cognitive Load and Autonomic Arousal During Voice Production and Vocal Auditory-Motor Adaptation. J Speech Lang Hear Res. 2025;68(4):1634–53.

57. Mesagno C, Hill DM. Definition of choking in sport: Re-conceptualization and debate. International Journal of Sport Psychology. 2013;44(4):267–77.

58. McNevin NH, Shea CH, Wulf G. Increasing the distance of an external focus of attention enhances learning. Psychol Res. 2003;67(1):22–9.

59. Beilock SL, Kulp CA, Holt LE, Carr TH. More on the fragility of performance: choking under pressure in mathematical problem solving. J Exp Psychol Gen. 2004;133(4):584–600.

60. Wulf G, Prinz W. Directing attention to movement effects enhances learning: A review. Psychonomic Bulletin & Review. 2001;8(4):648–60.

61. Wulf G, McNevin N, Shea CH. The automaticity of complex motor skill learning as a function of attentional focus. Q J Exp Psychol A. 2001;54(4):1143–54.

62. Wulf G, Hoss M, Prinz W. Instructions for motor learning: differential effects of internal versus external focus of attention. J Mot Behav. 1998;30(2):169–79.

63. Wulf G, Lauterbach B, Toole T. The learning advantages of an external focus of attention in golf. Res Q Exerc Sport. 1999;70(2):120–6.

64. Wulf G, McConnel N, Gartner M, Schwarz A. Enhancing the learning of sport skills through external-focus feedback. J Mot Behav. 2002;34(2):171–82.

